# A framework integrating multiscale in-silico modeling and experimental data predicts CD33CAR-NK cytotoxicity across target cell types

**DOI:** 10.1101/2024.12.31.630941

**Authors:** Saeed Ahmad, Kun Xing, Harshana Rajakaruna, William C. Stewart, Kyle A. Beckwith, Indrani Nayak, Meisam Naeimi Kararoudi, Dean A. Lee, Jayajit Das

## Abstract

Uncovering mechanisms and predicting tumor cell responses to CAR-NK cytotoxicity is essential for improving therapeutic efficacy. Currently, the complexity of these effector-target interactions and the donor-to-donor variations in NK cell receptor (NKR) repertoire require functional assays to be performed experimentally for each manufactured CAR-NK cell product and target combination. Here, we developed a computational mechanistic multiscale model which considers heterogenous expression of CARs, NKRs, adhesion receptors and their cognate ligands, signal transduction, and NK cell-target cell population kinetics. The model trained with quantitative flow cytometry and in vitro cytotoxicity data accurately predicts the short- and long-term cytotoxicity of CD33CAR-NK cells against leukemia cell lines across multiple CAR designs. Furthermore, using Pareto optimization we explored the effect of CAR proportion and NK cell signaling on the differential cytotoxicity of CD33CAR-NK cells to cancer and healthy cells. This model can be extended to predict CAR-NK cytotoxicity across many antigens and tumor targets.

## Introduction

Natural killer (NK) cells are lymphocytes of the innate immune system and interact with target cells using a wide array of activating, inhibitory, and adhesion receptors expressed on the cell surface (1). Integration of opposing signals initiated by diverse receptor-ligand interactions determine cytotoxic or tolerogenic response of NK cells towards their targets (2). Tumor or virally infected cells upregulate ligands cognate to activating NK cell receptors (NKRs) or downregulate ligands cognate to inhibitory NKRs which usually lead to lysis of the tumor cells by interacting NK cells (3). NK cells may be engineered to express chimeric antigen receptors (CARs) that expand their repertoire of recognized antigens (4–6). Compared to CAR-T cell based therapies, CAR-NK cell therapy potentially minimizes both antigen escape and off-tumor toxicities by avoiding reliance on the expression of a single targeted antigen and utilizing healthy tissue-sparing inhibitory signaling (7).

Understanding and predicting tumor cell responses to CAR-NK cytotoxicity is essential for designing approaches to improve therapeutic efficacy. The overall anti-tumor activity of each manufactured CAR-NK product is variable and dependent on many factors including the efficiency of CAR transduction, tumor expression of the targeted antigen, and the innate cytolytic capacity of the non-modified NK cell starting material. Due to this complexity, the therapeutic efficacy of each manufactured CAR-NK cell product should be assessed individually through time and resource consuming functional assays. Computational modeling offers an avenue to explore the biological parameters which impact CAR-NK cytotoxicity and predict real-world outcomes.

Several previous computational models have explored the CAR design space, primarily in the context of CAR-T cells. These have revealed important parameters such as antigen-receptor affinity tuning, dosing protocols, and cellular transition states that predict biological outcomes and, ultimately, patient responses to CAR immunotherapies (8–13). However, computational modeling of CAR-NK cells presents a unique challenge compared to modeling CAR-T cells due to the diverse array of germline-encoded activating and inhibitory receptors that regulate their innate cellular cytotoxicity (2). Since CAR-NK cell effector functions are determined by the integration of signals initiated by both the CAR and the diverse NKRs and adhesion receptors, the cytotoxic response of CAR-NK cells may not follow an intuitive dependence on the availability of the CAR antigens or strength of signaling domains. Accounting for this diversity of receptors is challenging computationally as the decision-making process of NK cells cytotoxicity is still incompletely understood, despite insights from mathematical modeling (14, 15). Indeed, a previous agent-based model of CAR-NK cell cytotoxicity does not take into account the expression of co-receptors at the single-cell level that are responsible for NK cell’s innate cytotoxicity (16).

Here we develop a framework combining *in vitro* experiments with a multi-scale mechanistic *in-silico* model which integrates single-cell level ligand and receptor expressions, single cell signal transduction, and population kinetics of CAR-NK cells and target cells to quantitatively explore the roles of CAR, activating and inhibitory NKRs, and adhesion receptors in determining cytotoxicity against tumor and healthy cells. Our multiscale *in-silico* model is trained with single-cell abundances of tumor ligands and NKRs measured by quantitative flow cytometry along with *in vitro* cytotoxicity data. Our model predicts short- and long-term CAR-NK cell cytotoxicity against novel tumor cell lines and healthy cells. Furthermore, we have employed a Pareto optimization approach to determine the optimal CAR expression and NK cell signaling which enables CAR-NK cell to discriminate between tumor cells and healthy cells. This model serves as a proof-of-concept tool for designing, evaluating, and predicting CAR-NK cell cytotoxicity and can be adapted and trained for a wide range of predictive clinical and translational applications.

## Results

### CAR-NK cytotoxicity displays nonlinearity with antigen expression

It has been shown experimentally that CAR-T cell activation increases monotonically with increasing cognate antigen expression (Fig. 1A) (17, 18). This is reasonable as the increased expression of antigens on target cells give rise to increased CAR-antigen complexes, increasing the strength of the CAR signaling, and consequently increasing CAR T cell effector functions such as cytotoxicity. Prior mathematical models of CAR-T cell cytotoxicity based on this reasoning are consistent with experimental outcomes (9, 10, 19, 20). To determine whether CAR antigen density alone would account for the dynamic variability in CD33CAR-NK cytotoxicity, we performed a flow cytometry experiment and correlated CD33 expression with CD33CAR- NK cytotoxicity. We observed a non-monotonic relationship between antigen density and cytotoxicity across multiple donors and CAR designs, indicating that modeling NK cell activation through CAR antigen expression alone is not sufficient to account for the observed cytotoxicity (Fig. 1B). We observed similar non-monotonicity of CAR-NK cell cytotoxicity with both adhesion (ICAM-1) and inhibitory (HLA-ABC) ligand expression on target cells (Fig. S1). Thus, we reasoned that a model which considers multiple ligand and receptor interactions would be necessary to predict CAR-NK cell cytotoxicity.

**Figure 1:**
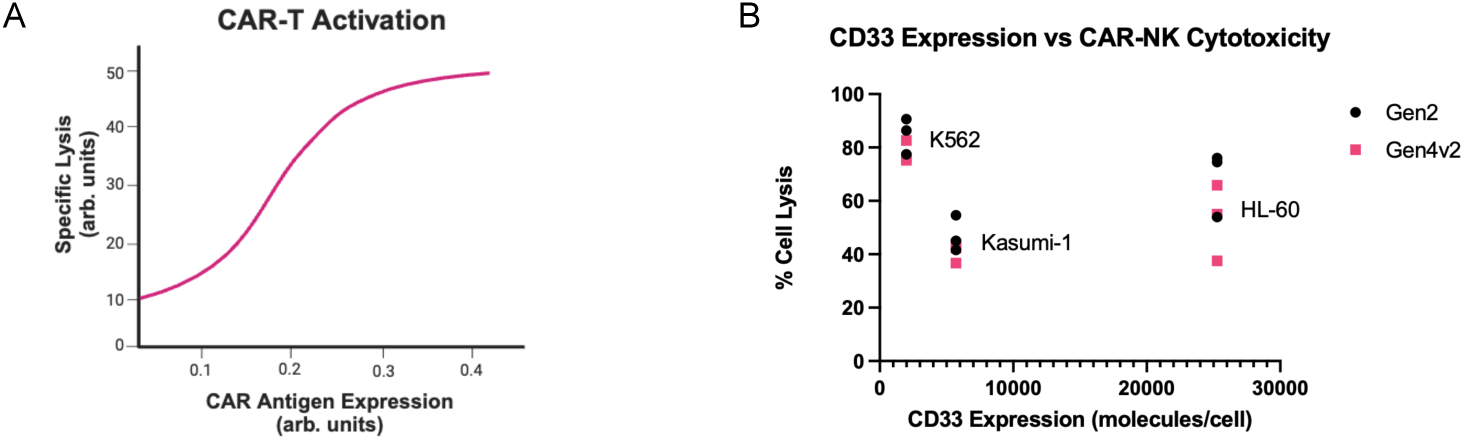
CD33CAR-NK cytotoxicity displays non-monotonicity with CD33 expression. A. Representative monotonic relationship between CAR-T activation and CAR antigen expression, extrapolated from RA Hernandez-Lopez 2021. B. Non-monotonic relationship between measured CD33 expression on K562, Kasumi-1, and HL-60 and CAR-NK cytotoxicity against, K562, Kasumi-1, and HL-60 cell lines. N = 3 donors.

### Model predicts killing of HL-60 by CD33CAR-NK in a donor specific manner

We developed a multiscale mechanistic *in-silico* model which integrates signals arising from interaction of CARs, inhibitory, activating, and adhesion NK cell receptors (NKRs) with their cognate ligands expressed on target cells and predicts the cytotoxicity of CD33CAR-NK cells against target cells. In this model, we include key processes involving interaction of NK cells with target cells (e.g., tumor or healthy cells) through processes that occur in three different scales: molecular, sub-cellular, and cell population. *Molecular scale*: In the model, CD33CAR, adhesion receptor LFA-1, and inhibitory KIRs bind their cognate ligands CD33, ICAM-1, and HLA-ABC respectively. These interactions are described by second-order binding-unbinding reactions with their respective binding and unbinding rates. The effects of activating NKRs are included implicitly (details in Materials and Methods). The CD33CAR constructs (Gen2, Gen4v2, V5, and V6) we considered here contain CD3ζ and other activating domains such as 4- 1BB, CD28, and 2B4 (Fig. 2D). We account for the cell-cell variations of receptors and their cognate ligands in the model by considering the distribution of their expressions instead of their average value. *Subcellular scale*: The formation of the ligand-receptor complexes initiates different downstream reactions within the NK cell (21). Stimulatory proteins such as CD3ζ in the intracellular domain of the CARs are phosphorylated by Src family kinases leading to recruitment of Syk family kinases (22). The phosphorylated Syk family kinases induce further downstream signaling reactions leading to phosphorylation of the guanine exchange factor Vav1. Phosphorylated Vav1 leads to the release of lytic granules onto the target cells (23, 24). Upon binding with ligand ICAM-1, the adhesion receptor LFA-1, promotes phosphorylation of Vav1 (25). Thus, both CAR and the adhesion receptor contribute toward Vav1 phosphorylation. The inhibitory KIRs are associated with immunoreceptor tyrosine-based inhibition motifs (ITIMs) in the transmembrane domain which are phosphorylated by Src family kinases upon binding with class I MHC molecules. Phosphorylated ITIMs recruit the phosphatase SHP-1 which deactivates Vav1 via dephosphorylation (26). We used a series of first order reactions to represent the above chemical modifications as different ‘states’ of the receptor ligand complex (Fig. 2B and Methods). The CAR, adhesion, and the inhibitory KIR signaling lead to formation of the end complexes, C_N1_, C_N3_, and C_N4_, respectively (Fig. 2B). We did not model the activating NKR signaling explicitly as there are multiple activating NKRs in the donor NK cells which could be stimulated by uncharacterized ligands on target cells. Therefore, we introduced the end complex corresponding to all other activating receptor signaling as the parameter C_N2_. In the model, the end complexes C_N1_, C_N2_, and C_N3_, phosphorylate Vav1 as enzymes. The end complex C_N4_ corresponds to inhibitory KIR signaling dephosphorylates phosphorylated Vav1 enzymatically. The details of the model are provided in the Materials and Methods. *Cell population*: We consider a population of target and NK cells interacting where each interaction between a NK cell and a target cell leads to production of phosphorylated Vav1 and subsequent lysis of the target cell with a rate proportional to the abundance of phosphorylated Vav1 generated during the interaction. We also assume each NK cell can interact with any target cell with equal probability. The tumor target cells can proliferate however we assume the NK cells do not proliferate or die within the time scale of interest (∼4 to ∼48 hours). The population kinetics of target and NK cells are given by a set of coupled ordinary differential equations (ODEs) (See Materials and Methods). The *in-silico* model contains nine parameters *α*_1_, *C_N2_*, *α*_3_, *α*_4_, *V_c_*, *K*_1_, *K*_2_, *λ_c_*, *r_c_* that are estimated during model training using single cell abundances of the receptors and ligands, and percentage of lysed target cells in *in vitro* cytotoxicity assays. The list of the parameters along with their interpretations are shown in Table 1.

**Figure 2.**
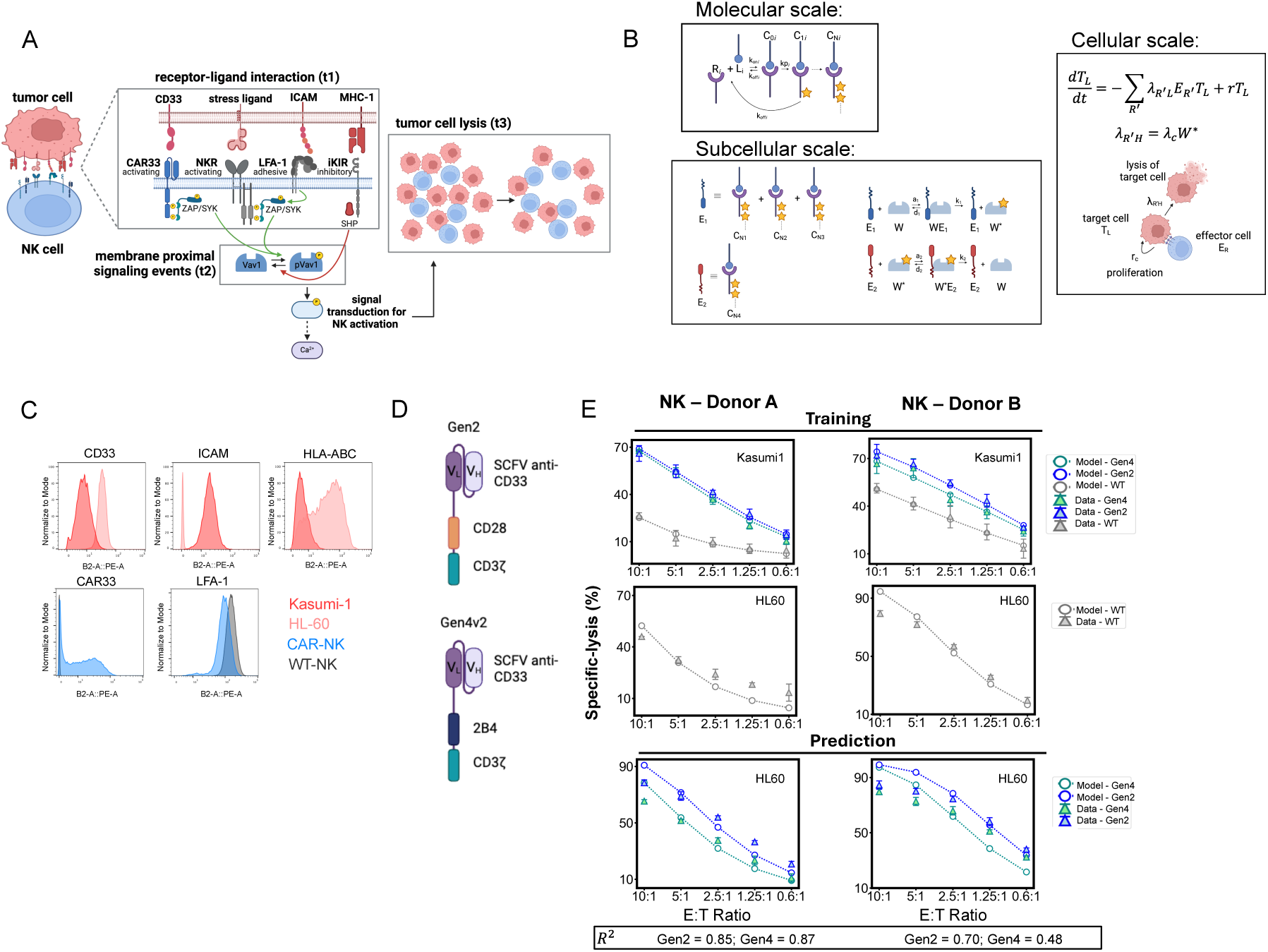
Multiscale in-silico model design and cytotoxicity prediction of CAR-NK tumor lysis against HL-60. A. Representation of CD33CAR-NK activation and cytotoxicity shown via three biological steps. B. Representative schematics of modeled reactions. C. (Top) Expressions of CD33, ICAM, HLA-ABC on Kasumi-1 and HL60. (Bottom) CD33CAR and LFA-1 expression on CAR-NK and WT-NK cells. D. CAR designs evaluated in A and E. E. (Top and middle) Training data and model fit with CD33CAR-NK and WT-NK cytotoxicity against Kasumi-1 and WT-NK cytotoxicity against HL-60. (Bottom) Prediction of cytotoxicity against HL-60. Cytotoxicity was performed with calcein for four hours. R^2^ coefficient of determination calculated for goodness of fit with the data.

**Table 1:** List of processes and parameter values used in the model. * denotes fixed values.

|  | Parameter<br>(unit) | Description | Units | Parameter<br>range | Kasumi1-DonorA<br>Estimate<br>(CI) | Kasumi1-DonorB<br>Estimate<br>(CI) |
| --- | --- | --- | --- | --- | --- | --- |
| 1 | $\alpha_1(\text{Ge4})$<br>(dimensionless) | forward probability of active complex of CD33CAR and CD33 | Unitless | 0-1 | 0.74<br>(0.35-1.0) | 0.51<br>(0.10-0.92) |
| | $\alpha_1(\text{Gen2})$ | forward probability of active complex of CD33CAR and CD33 | Unitless | 0-1 | 0.68<br>(0.32-1.0) | 0.38<br>(0.07-0.70) |
| 2 | $C_{2N}$ | concentration of activating receptor-ligand active complex | Molecules/cell | 100-5000 | 608.10<br>(100-2165) | 1234.15<br>(100-2967.8) |
| 3 | $\alpha_3$ | forward probability of active complex of LFA-1 and ICAM-1 | Unitless | 0-1 | 0.20<br>(0.04-0.35) | 0.48<br>(0.07-0.90) |
| 4 | $\alpha_4$ | forward probability of active complex of iKIR and HLA-ABC | Unitless | 0-1 | 0.71<br>(0.46-0.97) | 0.68<br>(0.32-1.0) |
| 5 | $V_c$ | Relative rate of pVav1 formation | Unitless | $10^{-3}$ to $10^{-1}$ | 0.27<br>(0.11-0.43) | 0.28<br>(0.02-0.53) |
| 6 | $K_1$ | Scaled Michaelis constant for transition of Vav1 to pVav1 | Unitless | 0.01 to 1 | 0.001<br>(*) | 0.09<br>(0.001-0.56) |
| 7 | $K_2$ | Scaled Michaelis constant for transition of pVav1 to Vav1 | Unitless | 0.01 to 1 | 0.052<br>(0.033,0.072) | 0.002<br>(0.001-0.008) |
| 8 | $\lambda_c$ | Lysis constant | $hr^{-1}$ | $10^{-4}$ to $10^{-7}$ | 2.0e-5<br>(1.7e-5,2.3e-5) | 4.2e-5<br>(2.5e-5-6.0e-5) |
| 9 | $r_c$ | Proliferation rate of tumor cells | $hr^{-1}$ | 0 to $10^{-3}$ | 0 | 0 |

We trained the above multiscale *in-silico* model and then evaluated the model’s prediction of short term (4 hours) cytotoxicity of CD33CAR-NK cells to novel cell lines. We performed quantitative flow cytometry to approximate the molecular abundances of CD33, ICAM-1, and HLA-ABC on AML cell lines Kasumi-1 and HL-60 as well as CD33CAR and LFA-1 on NK cells (Fig. 2C, Fig. S2). KIR abundances were taken from reported values (27). We predicted the cytotoxicity of two CAR-NK designs, Gen2 and Gen4v2 which differ in their signaling domains (6). We split cytotoxicity data into train and test data sets and trained the model by first fitting the lysis of WT-NK and CAR-NK cells against Kasumi-1 to the eight parameters *α*_1_, *C*_𝑁2_, *α*_3_, *α*_4_, *V*_c_, *K*_1_, *K*_2_, *λ*_c_, *r*_c_. As the doubling time for AML cell lines is between one to two days, *r*, the parameter reflecting the proliferation rate of target cells, was set to 0 for models trained on short-term (4 hr) cytotoxicity assays (28). Using the common parameter values identified, we then re-estimated *C*_𝑁2_,and *λ*_c_, for HL-60 to account for the differences in the activating ligand repertoire and lysis sensitivity of HL-60. Parameter estimation of *α*_1_, *C*_𝑁2_, *α*_3_, *α*_4_, *V*_c_, *K*_1_, *K*_2_, *λ*_c_ was performed through minimizing the least square residual cost function (See Methods). The estimated values of the parameters and the confidence intervals are listed in Table 1. We set up an *in-silico* cytotoxicity assay with a mixture of 10,000 target cells, varying the number of effector cells to recapitulate the E:T ratios used experimentally. The NK cells and target cells interacted in the model using the previously identified best fit parameters (Table 1). We then predicted the percentage of HL-60 tumor cells lysed by Gen2 and Gen4v2 CAR-NK after 4 hr of co-culture. We calculated goodness of fit of our model prediction using the 𝑅^2^ coefficient of determination and observed that the predictions agreed with the data for both Gen2 (𝑅^2^: 0.85, 0.70) and Gen4v2 (𝑅^2^: 0.87, 0.48) CAR designs in two NK cell donors (Fig. 2E). These results show that our multiscale *in-silico* model accurately predicts short term (4 hr) CD33CAR-NK cell lysis of novel tumor cells.

### Model predicts long term cytotoxicity of CD33CAR-NK cells

We next evaluated the ability of our model to predict the long-term (∼48-72 hr) cytotoxicity of CAR-NK cells. Long-term cytotoxicity assays spanning multiple days better capture CAR-NK cell serial killing (29). NK cell cytotoxicity mechanisms captured in long-term cytotoxicity assays include perforin/granzyme mediated lysis, secretion of soluble factors such as 𝐼𝐹𝑁𝛾, and death receptor interactions such as Fas-FasL (30). Although perforin-granzyme mediated cytotoxicity is the dominant signal encoded by the CAR and is the major mechanism of NK cell cytotoxicity, the secondary effects of cytokine secretion and antigen-independent death receptor signaling also play an important role in tumor control (31, 32). It was unknown how well the model would capture these additional functions of NK cells in a long-term cytotoxicity assay. To adapt the model to predict long-term CAR-NK cytotoxicity, we added one additional parameter, *r*_c_, to represent the proliferation rate of tumor cells. To assess the long-term killing ability of CAR-NK cells, we performed cytotoxicity assays with CD33CAR-NK cells and MV4-11 tumor cells and measured target cell death after 48 and 72 hr of coculture using propidium iodide flow cytometry. We evaluated the cytotoxicity of two CD33CAR constructs, V5 and V6, which differ in their orientation of the light and heavy chains of the scFv, which could impact ligand affinity, but are otherwise identical in all other domains (Fig. 3B). We split the cytotoxicity data into test and train data sets and trained the model using CD33CAR-NK and WT NK cytotoxicity of MV4-11 target cells at 48 hr. Molecular abundances of the modeled receptor and ligands were approximated using quantitative flow cytometry (Fig. 3A and Fig. S2A). After fitting the model to the data and parameter estimation, we set up an *in-silico* cytotoxicity assay and predicted the percent lysis of MV4-11 cells by CD33CAR-NK cells and WT-NK cells at 72 hrs. We observed good agreement between the *in silico* model prediction and the data (V5 𝑅^2^: 0.87, 0.96, 0.99 and V6 𝑅^2^: 0.99, 0.95, 0.87, WT 𝑅^2^: 0.57, 0.77, 0.51) from *in vitro* experiments of three independent donors, showing that our multi-scale model robustly captures NK serial killing and multiple mechanisms of tumor cytotoxicity (Fig. 3C).

**Figure 3.**
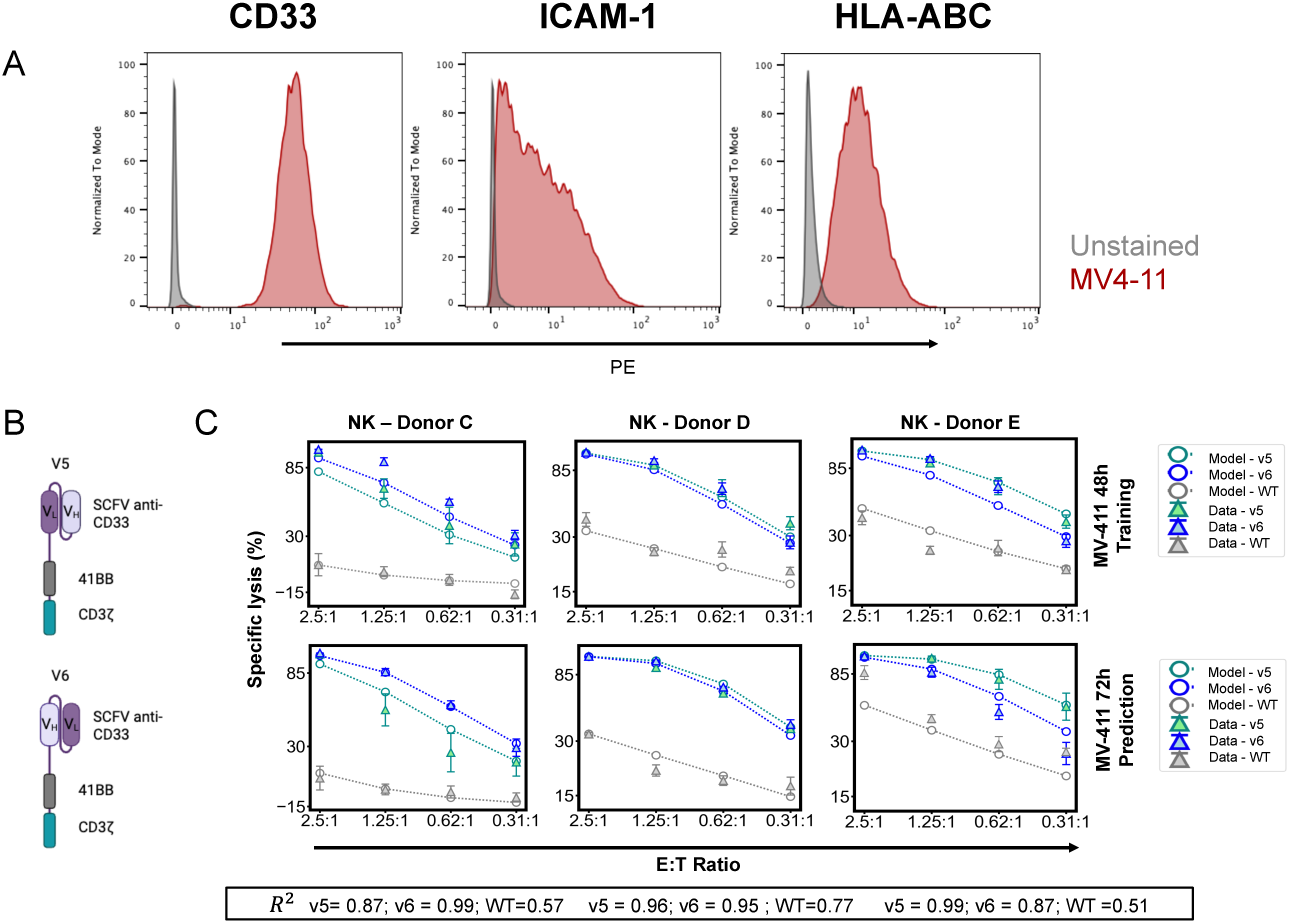
Model predicts long-term CD33CAR-NK cytotoxicity against MV4-11. A. Distribution of CD33, ICAM-1, and HLA-ABC expression on MV4-11 tumor cells by flow cytometry. B. CAR designs used in cytotoxicity assays and evaluated in C. (Top) Training data and model fit with CD33CAR-NK and WT-NK cytotoxicity against MV4-11 at 48 hr. (Bottom) Model prediction of CD33CAR-NK and WT-NK cytotoxicity against MV4-11 at 72 hr. Cytotoxicity performed with propidium iodide flow cytometry. N = 3 donors.

### CD33CAR expression results in targeting of healthy monocytes

On-target off-tumor toxicities arise when CAR expressing immune cells target non-malignant tissues that express the targeted antigen (33). In many cancers, including acute myeloid leukemia (AML), there is a lack of ideal antigens that are exclusively expressed on cancer cells and absent from healthy cells. For example, CAR therapies designed to target myeloid antigens such as CD33, CD123, and CD38 inadvertently result in cytotoxicity against healthy myeloid cells which express low to moderate amounts of the cognate antigen (34, 35). Mis-targeting of CD33, specifically, results in lysis of healthy cells such as neutrophils and monocytes which are essential for controlling infection (36, 37). These on-target off-tumor toxicities have posed significant clinical risk in CAR-T therapies (38, 39). The toxicities of CAR-NK cell therapies, including those associated with on-target off-tumor activity, have been less severe than those observed in CAR-T cell therapies, possibly due to the activation of inhibitory KIRs by MHC molecules (40, 41). However, these studies have been done in systems where the targeted antigen exhibits significantly lower expression on healthy tissues than the targeted tumor cell (41, 42). Similar on vs off tumor studies for targeted antigens which have equivalent expression on healthy and tumor cells have not been performed for CAR-NK cells. To evaluate the on-tumor vs off-tumor activity of CD33CAR-NK cells, we isolated primary monocytes from healthy donor PBMCs as a representative cell type of the myeloid lineage. We quantified ligand expression on monocytes using quantitative flow cytometry and found comparable levels of CD33 expression on monocytes and AML cell lines (Fig. 4A). One day after isolation, we performed calcein cytotoxicity assays using CD33CAR-NK cells or WT-NK cells without prior HLA matching. We observed that WT NK cells had low (4-25% lysis) cytotoxic activity against monocytes at all effector-to-target ratios (Fig. 4B). CD33CAR-NK cells showed improved cytotoxicity against Kasumi-1 tumor cells but also markedly increased cytotoxicity against healthy monocytes despite high levels of MHC Class I expression (Fig. 4C).

**Figure 4.**
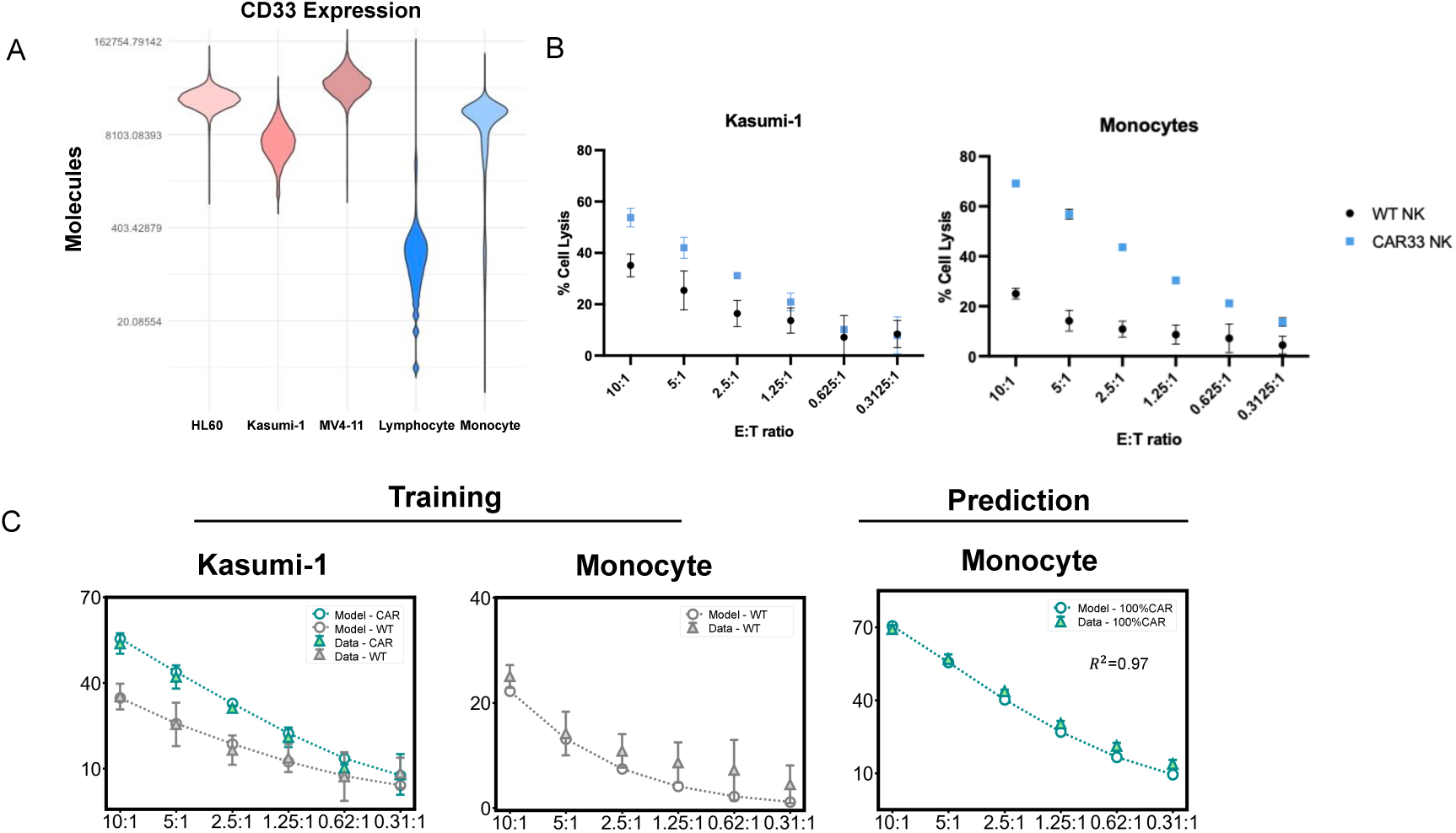
CD33CAR expression results in targeting of healthy monocytes. A. Expression of CD33 on monocytes, lymphocytes, and AML tumor cell lines by quantitative flow cytometry. B. calcein cytotoxicity assay of CD33CAR-NK and WT-NK cells against Kasumi-1 cells and primary monocytes. C. (Left) Training data and model fit of CD33CAR-NK and WT-NK cytotoxicity against Kasumi-1 and (Middle) WT-NK cytotoxicity against monocytes. (Right) Prediction of CD33CAR-NK cytotoxicity against primary monocytes. Cytotoxicity performed with calcein for four hours. *R2* coefficient of determination calculated for goodness of fit with the data.

Given these results, we first asked whether the model could predict the level of monocyte lysis observed in our experimental data. We employed the aforementioned workflow and trained the model using CD33CAR-NK and WT-NK cytotoxicity data against Kasumi-1 targets along with WT-NK cytotoxicity against monocytes from an un-matched donor. We trained the model to the data by estimating the parameters described (Table 1) and predicted the cytotoxicity of CD33CAR-NK cells to monocytes. We observed excellent agreement between our model prediction and the experimental data (𝑅^2^: 0.97), indicating that the modeled parameters and processes could be extended to describe NK cytolytic activity to non-malignant primary cells as well as tumor cells.

### Predicting optimal CAR expression and signaling to minimize on-target off-tumor cytotoxicity

After validating the model’s ability to describe non-tumor cytotoxicity, we aimed to manipulate parameters within the model framework to predict the optimal CAR expression and signaling kinetics to maximize cancer cell killing while sparing lysis of healthy tissues. In our experimental cytotoxicity data, we observed a tradeoff between cancer cell lysis and monocyte lysis (Fig. 5A) which suggested the system can be suitable for multi-objective parameter optimization. In multi-objective optimization, there are multiple desired outcomes which must be simultaneously considered, for example, maximizing lysis of cancer cells while minimizing lysis of healthy cells. Multi-objective optimization will produce a set of solutions which lie on a front known as the Pareto optimal front on which all points are equivalently optimal. Our data suggested that WT NK cells best minimized on-target off-tumor cytotoxicity while maintaining tumor-specific lysis, but it was unclear whether WT-NK cells represented the Pareto optimal front and which signaling parameters produced this front (Fig. 5A). We employed an *in-silico* two-objective Pareto optimization scheme to explore the signaling parameters of CAR-NK and WT-NK cells and calculate the Pareto optimal front, beyond which either lysis of cancer cells cannot be maximized and lysis of healthy cells cannot be minimized without sacrificing the other. In our two-objective optimization problem, we simultaneously minimized lysis of monocytes while maximizing lysis of Kasumi-1 tumor cells using *in-silico* cytotoxicity assays (43, 44). Effector NK cells were introduced at T=0 and allowed to interact with target cells for 4 hr while varying parameters to evaluate the Pareto fronts.

**Figure 5.**
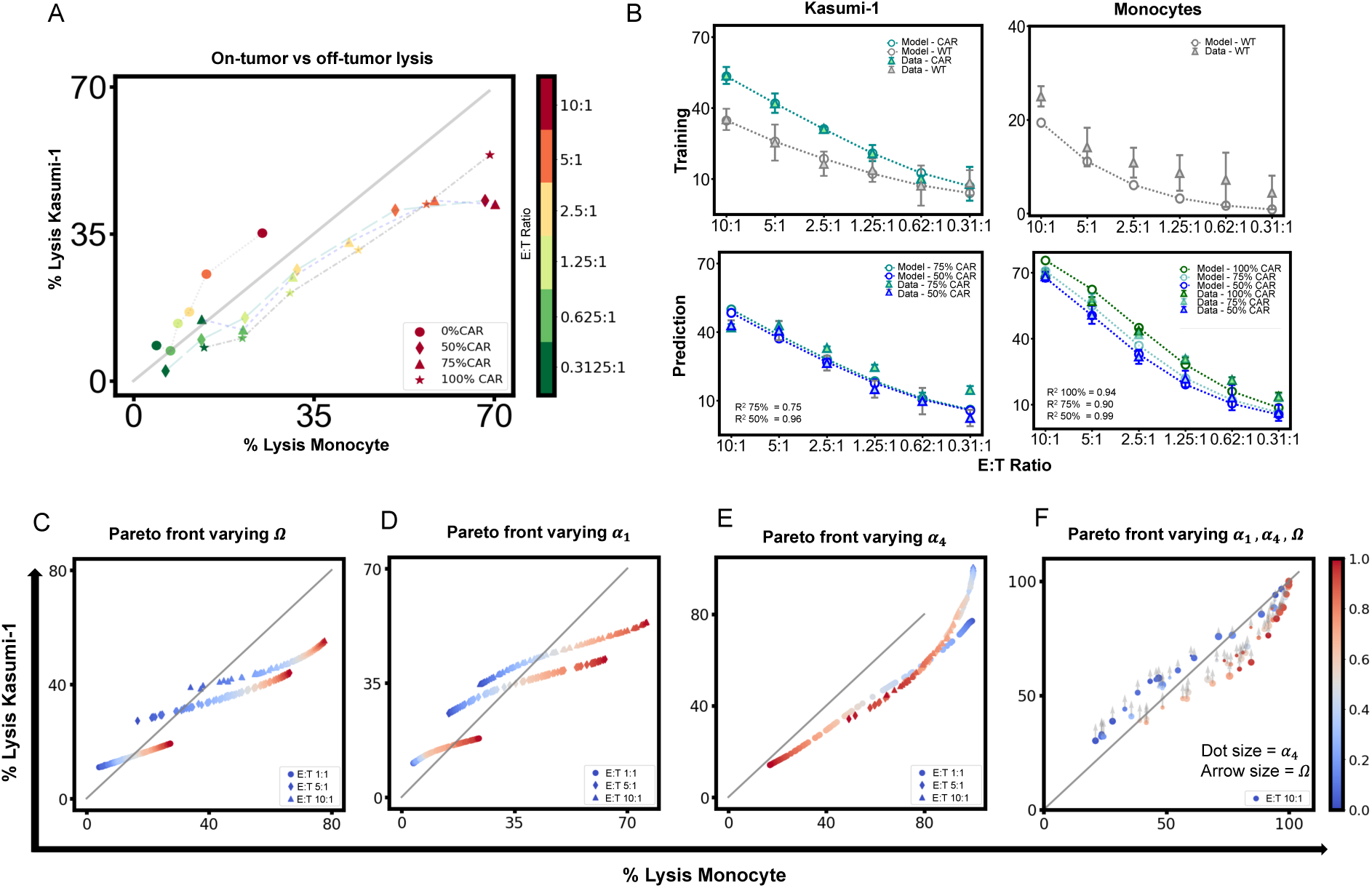
Pareto optimization reveals optimal responses of WT and CD33CAR NK cells against leukemic and healthy cells. A. On-tumor vs off-tumor cytotoxicity of CAR-NK and WT-NK cells. Line denotes % lysis of monocytes = %l lysis of Kasumi-1. B. (Top) Model fit of CD33CAR-NK and WT-NK cytotoxicity against Kasumi-1 and WT-NK cytotoxicity against monocytes. (Bottom) Prediction of 3:1 CD33CAR-NK:WT-NK cytotoxicity and 1:1 CD33CAR- NK:WT-NK against Kasumi-1 and monocytes. Mixtures created using estimated Ω values. *R2* coefficient of determination calculated for goodness of fit with the data. C-F: Pareto fronts varying different controlled parameters. Color scale bar refers to parameter varied in each figure. C. Pareto fronts for CAR-NK % lysis of monocytes and % lysis of Kasumi-1. The Pareto fronts are calculated for different E:T ratios, varying Ω as the controlled parameter. All other parameters are fixed to estimated values. D. Pareto fronts for CAR-NK % lysis of monocytes and % lysis of Kasumi-1. The Pareto fronts are calculated for different E:T ratios, varying *α*_1_ as the controlled parameter. All other parameters are fixed to estimated values. E. Pareto fronts for CAR-NK % lysis of monocytes and % lysis of Kasumi-1. The Pareto fronts are calculated for different E:T ratios, varying *α*_4_as the controlled parameter. All other parameters are fixed to estimated values. F. Pareto fronts for CAR-NK % lysis of monocytes and % lysis of Kasumi-1. The Pareto fronts are calculated for different E:T ratios, varying *α*_1_, *α*_4_, and Ω as the controlled parameters. *α*_1_ value given by color scale, *α*_4_ value given by dot size, Ω given by arrow size.

Firstly, we aimed to analyze the effect of CAR expression in minimizing off-tumor toxicity while maintaining on-tumor killing. The CD33CAR expression on CAR-NK cells showed a bimodal distribution which can be modeled as a mixture of two log-normal distributions with modes centered around low (0.42 MFI ≡467 molecules) and high (16.4 MFI ≡17188 molecules) MFI values (Fig. S3A). We created a parameter Ω which determines the percentage of CD33CAR positive NK cells and is assumed to be biologically equivalent to the transduction efficiency of CAR generation via CRISPR-AAV site-directed insertion. Given the presence of background cellular autofluorescence and non-specific antibody binding, we reasoned that the log-normal distribution centered around the lower MFI in the CAR-NK cells is equivalent to non-transduced WT-NK cells. Similarly, we reasoned that the lognormal distribution centered around the higher MFI represents the CAR expression in transduced NK cells. We modeled the cell surface expression of CD33CAR on CAR-NK cells and WT-NK cells as a mixture of two lognormal distributions and estimated the distribution parameters including Ω using maximum likelihood estimation (Fig. S3A, See Materials and Methods).

Next, we evaluated the ability of our *in-silico* model to describe and predict target cell lysis generated by the mixture of WT and CAR NK cells. We considered the following ratios of WT and CAR-NK mixtures (CD33CAR-NK alone, 3:1 CD33CAR-NK:WT NK, 1:1 CD33CAR-NK:WT-NK, and WT-NK alone) and measured cytotoxic responses of those mixtures against healthy and tumor target cells. We determined the value of Ω in the experimental mixtures through maximum likelihood estimation (Fig. S3A). We performed *in-silico* cytotoxicity assays using 10,000 target cells while varying Ω. Our model predictions varying Ω showed strong agreement with the biological data using both Kasumi-1 (3:1 CD33CAR-NK:WT NK: Ω = 0.55, 𝑅^2^ = 0.75; 1:1 CD33CAR-NK:WT-NK Ω = 0.44, 𝑅^2^: 0.96) and monocytes (CD33CAR-NK: Ω = 0.80, 𝑅^2^ = 0.94; 3: 1 CD33CAR − NK: WT NK: Ω = 0.55, 𝑅^2^: 0.90; 1:1 CD33CAR-NK:WT-NK: Ω = 0.44, 𝑅^2^ = 0.99) as the target cell for all NK cell conditions (Fig. 5B). Similarly, the model was also able to predict cytotoxicity generated by a mixture of cells when the mixture of effector cells was generated by sampling rather than Ω estimation (Fig. S3B). Therefore, the *in-silico* model can successfully describe target cell lysis by CD33CAR-NK cells with different CAR expression efficiency.

We then computationally investigated the effect of manipulating transduction efficiency, or Ω, on the trade-off between healthy cell and tumor cell lysis through performing *in-silico* cytotoxicity assays while varying Ω. Concordant with our experimental data, we found that increasing Ω increased lysis of Kasumi-1 cells but also disproportionally increased lysis of healthy monocytes (Fig. 5C).

Next, we performed Pareto optimization while varying signaling parameters *α*_1_and *α*_4_to investigate the signaling mechanisms which produce the Pareto optimal front. *α*_1_ represents the signaling propensity of CD33CAR and we found that increasing *α*_1_increased lysis of Kasumi-1 cells but disproportionally affected lysis of healthy monocytes (Fig. 5D). Decreasing *α*_4_, which represents the signaling propensity of inhibitory receptors, similarly increases lysis of cancer cells and healthy cells. The differential effect of *α*_4_ on lysis is E:T ratio dependent, where for low E:T ratios, changing *α*_4_ has more impact on healthy cell lysis than cancer cell lysis (Fig. 5E). Simultaneously varying *α*_1_, *α*_4_, 𝑎𝑛𝑑 Ω revealed that NK cells which are representative of WT-NK cells (i.e. cells with either low CAR transduction or low CAR signaling) best optimized the objective, confirming that WT-NK cells represent the Pareto optimal front (Fig. 5G).

## Discussion

We developed a framework integrating quantitative flow cytometry measurements of receptor and ligand expressions and *in vitro* cytotoxicity assays with multi-scale mechanistic computational modeling that predicts WT and CD33CAR-NK cell cytotoxicity across different tumor cell lines. We utilized a coarse-grained modeling approach which implicitly includes subcellular signaling processes into the model parameters, allowing us to focus our quantitative analysis on select receptor and ligand abundances and signaling parameters which are important for CAR-NK cytotoxicity.

The *in-silico* model integrates opposing signaling initiated by CAR, activating NKRs, adhesion receptors, and inhibitory NKRs and correctly captures the non-monotonicity of CAR-NK cell cytotoxicity against tumor cells expressing various molecules of CAR antigens (e.g., CD33). The model also successfully describes cytotoxicity of CAR NK cells containing different CAR signaling domains such as CD28 in Gen2 CAR replaced by 2B4 in Gen 4v2 CAR, as well as short- (4 hr) and long-term (48-72 hr) cytotoxicity of the CAR NK cells. In addition, the model also correctly predicts cytotoxic response of WT NK cells and the mixture of WT and CAR NK cells. The ability of our model to describe lysis data and predict cytotoxicity outside of the training data with high accuracy shows the validity of our integrated framework. The framework can be straightforwardly extended to describe cytotoxicity of other CAR constructs and target cells. Together, these studies shows that CAR-NK cytotoxicity can be accurately modeled with a limited number of experimentally measurable variables and a conservative system of ordinary differential equations.

The multiscale model implicitly describes the subcellular signaling processes including immunological synapse formation (45) or inside-out LFA1-ICAM signaling (46) and cytotoxic responses using coarse-grained processes. Values of the associated model parameters can potentially capture the changes in the CAR transmembrane domain or any involvement of FasL-Fas interactions in late-time cytotoxicity. The model parameters account for unknown contributors to cytotoxicity such as intrinsic tumor resistance mechanisms against NK cytotoxicity (47, 48). Poor model predictions were largely due to saturations in percentage lysis in *in vitro* cytotoxicity assays where the experimentally measured cytotoxicity approached or exceeded 100% in training data (Fig. 2E). As an extension of this work, investigating poor model predictions may uncover nondominant mechanisms or discovering unknown mechanisms of CAR-NK cytotoxicity relevant to therapeutic efficacy.

We observed that per cell molecules of CAR and LFA vary over 100-fold and 10-fold in CAR NK cells, respectively, and the per cell molecules of CD33, ICAM1, HLA-A/B/C vary over 10- fold in the tumor target cells (HL60, AML10) and healthy monocytes. The cell-cell variability in receptor and ligand expression can also give rise to cell-cell variations of cytotoxic responses of CAR and WT NK cells. Our model captures these cell-cell variations of cytotoxicity arising due to variations of per cell molecules of CAR, activating and inhibitory NKRs, and adhesion receptors in NK cells and abundances of cognate ligands on target cells. The excellent agreement of the model predictions for the percentage lysis with the cytotoxicity assays suggest the relevance of such cell-cell heterogeneity in determining bulk cytotoxicity response. A model devoid of cell-cell variations in receptors and ligands and instead trained only by the mean values shows lower predictive capacity, particular in predicting lysis of healthy cells, (Fig. S2B). This finding supports the relevance of cell-cell heterogeneity in affecting the bulk cytotoxic response.

We applied our model to explore the on-tumor and off-tumor cytotoxic activity of CAR-NK cells. We performed multi-objective optimization to optimize the on-target cytotoxicity of CAR-NK cells. Our analysis manipulating parameters that reflect the efficiency of CAR transduction, CAR signaling strength, and inhibitory receptor signaling strength all converge and agree with the biologic data that WT-NK cells maximize on-tumor toxicity while minimizing off-tumor toxicity. This analytical approach is valuable for NK cell-based cancer immunotherapies where the cell and the CAR both contribute distinct and substantial anti-tumor cytotoxicity.

## Limitations of this study

Our model assumes limited genotypic and phenotypic polymorphisms in NK cell receptor repertoire and does not specify contributions of cytotoxicity from all known NK cell receptor-ligand interactions with tumor cells. This assumption is most directly reflected in the *C*_2𝑁_ parameter, which must be re-estimated for each additional target cell. We show that the granularity and number of the ligand-receptor pairs represented in the model is sufficient for a minimal and generalizable framework for predicting CAR-NK cytotoxicity. Although other identified receptor-ligand interactions play a role in NK cell cytotoxicity, our understanding of the hierarchy of signal integration as determinants of NK cytotoxicity is incomplete. Further work characterizing the NK-intrinsic and tumor-intrinsic factors that mediate NK cell cytotoxicity is necessary to develop more powerful predictive models.

Our model only takes in selected signaling kinetics and population dynamics parameters to predict cytotoxicity such as binding affinity, and the distribution of NK cells and tumor cells in a mixed co-culture. Experimentally, we did not directly measure NK cell activation and instead utilized cytotoxicity as a downstream surrogate for activation. As cytotoxicity is a combined function dependent on both effector activation and target response, there may be NK-intrinsic signaling overlooked by our approach and instead compensated by other parameters. To adapt this model to intricately study NK cell signaling, incorporating data which directly measures NK activation (ie. Ca2+ flux, signaling proteins) would be necessary. Furthermore, we have utilized only *in vitro* data in this model and thus do not consider CAR-NK transit, homing, exhaustion, or antigen escape in the model. As these parameters have been shown to be important for clinical responses to adoptive T and NK therapies, predictive *in-silico* models may also benefit from training on these types of data (49–51).

The model system used for Pareto optimization is not representative of all situations in which off-tumor toxicities would arise. In many cases of off-tumor toxicities, the targeted antigen is lowly or moderately expressed on healthy tissues and highly expressed on the targeted tumor, contrasting with the system of Kasumi-1 and primary monocytes we utilize in which the tumor cells have lower expression of CD33 (33). Pareto analysis using a tumor cell line (i.e., MV4-11) which abundantly expresses the targeted antigen and is resistant to innate WT-NK killing may reveal a Pareto optimal front that favors CAR-NK cells.

In this study, we have focused on CARs targeting CD33, a sialic acid receptor expressed on 90% of AML tumors and other cells of the myeloid lineage (52, 53). Unlike B cell targeting (ie. CD19) CARs which have more tolerable off-tumor toxicities due to readily available therapeutic interventions, CARs which target myeloid tumors have much more severe risks associated with off-tumor toxicities. The clinical problem of simultaneously targeting healthy tissues in CAR therapies is also pertinent in solid tumors. In the future, applying this model to evaluate the on-target off-tumor cytotoxicity of solid tumor-targeting CAR-NK cells is both exciting and clinically relevant.

## Materials and Methods

### Quantitative Flow Cytometry

Kasumi-1, HL-60, WT NK and CAR-NK cells were stained with Tonbo Ghost Dye (Tonbo Biosciences, San Diego, CA) in PBS and Fc blocked (Miltenyi 130-059-901) prior to staining with either anti-CD33 (BD 561816), anti-ICAM (BD 560971), anti-HLA-ABC (BD 567582), anti-CD11a (BD 550851), or anti-Whitlow linker (Cell Signaling 62405S) antibodies in flow buffer (PBS +2% FBS). Cell surface protein expression was quantified using PE phycoerythrin fluorescence quantitation kit according to manufacturer’s instructions (BD 340495).

### CAR-NK Generation and Expansion

CD33 CAR-NK were generated as described previously(6). Briefly, NK cells were isolated from PBMC buffy coats from healthy human donors using RosetteSep Human NK cell Enrichment Cocktail (STEMCELL Technologies, Vancouver, BC, Canada). Isolated NK cells were expanded with irradiated mbIL-21 and 4-1BBL expressing K562 feeder cells at a 2:1 ratio for one week prior to CAR generation as previously described (54). On day 7, NK cells were electroporated with Cas9-RNPs using HiFi SpCas9 Nuclease V3 (Integrated DNA Technologies, Coralville, IA) and gRNAs targeting AAVS1 or CD38. 30 minutes after electroporation, NK cells were transduced with AAV6 virus (Andelyn Biosciences) encoding the CD33 CAR construct at an MOI of 70K virions per cell. Two days after electroporation, cells stained for CAR expression and were expanded twice with irradiated mbIL-21 and 4-1BBL expressing K562 feeder cells at a 1:1 ratio.

### Monocyte Isolation

Primary monocytes were isolated from healthy donor buffy coats using RosetteSep Human Monocyte Enrichment Cocktail (STEMCELL Technologies, Vancouver, BC, Canada). Isolated monocytes were cultured for 24 hours in complete RPMI (RPMI + 10% FBS) media prior to use in calcein cytotoxicity assays.

### Flow Cytometry Based Cytotoxicity Assays

Target cells were washed in complete RPMI (RPMI + 10% FBS) and resuspended to a final concentration of 0.1e6 cells/mL. NK cells were resuspended in complete RPMI at a concentration of 1e6 cells/mL and serially diluted in a twofold dilution in a 96 well plate. Cells were plated in triplicate. 1e4 labeled target cells were added to each well of NK cells to achieve the final effector to target ratios 10:1, 5:1, 0.625:1, 0.3125:1. Target cells and NK cells were incubated for the allotted time at 37°C. After incubation, propidium iodide was added to each well and flow cytometric analysis was performed to identify dead target cells.

### Calcein Cytotoxicity Assays

Target cells were loaded with calcein AM (2ug/mL) ((Fisher Scientific, Hampton, NH).Target cells were incubated with serially diluted effector cells achieve the final effector to target ratios 10:1, 5:1, 0.625:1, 0.3125:1. Maximum release was achieved with media containing 2% Triton-X 100. Spontaneous release was achieved by incubating target cells with 100 uL of complete RPMI media. Cells were incubated at 37°C for 4 hours and supernatant was collected for reading on a fluorescence plate reader. Cytotoxicity is plotted as percent specific lysis [(experimental release− spontaneous release)/(maximum release−spontaneous release)]×100 (54).

### Solutions of the ODE model

In this model, we have assumed a simplistic scenario where the NK cells interact with the tumor cells across three different timescales (from fast to slow) as shown in Fig. 2A and 2B. In the first stage (at *Molecular scale*), we have considered the interaction between four types of NK receptors (NKRs): the CAR receptors CD33CAR, activating receptors (such as NKG2D), adhesive receptors (LFA1) and inhibitory receptors (KIR) to their cognate ligands expressed on tumor cells, namely CD33, stress ligands, ICAM, and HLA-ABC, respectively. These receptor-ligand interactions lead to formation of the activated complexes. In *the Subcellular regime,* the activated end complexes formed by binding of CD33CAR, activating NKRs, LFA1 with their cognate ligands act as kinases, that phosphorylate Vav1 to phosphorylated-Vav1 (pVav1).

Conversely, activated end complexes generated by KIR-HLA-ABC interaction behave as phosphatases that dephosphorylate pVav1 to Vav1. Our model assumes that the increased concentration of the pVav1 at *Cellular level* leads to the lysis of target cells by activating the NK cell.

Because the lysis of target cells occurs at much slower timescale (∼hours) than the rapid receptor signaling reactions (such as receptor-ligand interactions, interconversion of Vav1 and pVav1, which happen in order of seconds), we have correlated the steady-state conditions of the signaling species (here, steady state concentration of pVav1) with the lysis of the tumor cells.

*Molecular scale (Stage-I):* The cytotoxic process of NK cells which starts with binding of a receptor to its cognate ligand and culminated in the killing of a tumor cell involves numerous biochemical reactions within the signaling pathway. In our model, we have employed a kinetic proofreading scheme, wherein all the complexes or modified forms of signaling proteins (e.g., pVav1) generated in the signaling pathway become free and revert to their initial forms when the receptor unbinds from the cognate ligand (55).

The following is the signaling scheme of a specific type of receptor (R_i_) expressed on a NK cell interacting with its cognate ligand (L_i_) present on a tumor cell:

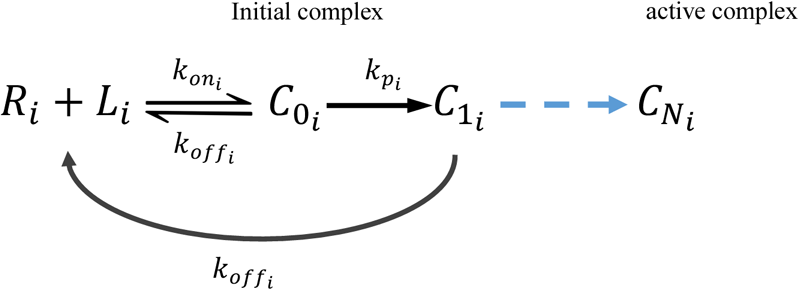

where index ‘*i*’ represents the types of the receptor 𝑅_𝑖_ (CD33CAR, activating NKRs, LFA-1, KIRs) that binds with its cognate ligand 𝐿_𝑖_ (CD33, stress ligands, ICAM, and MHC1) to form initial complex *C*_0𝑖_ with rate 𝑘_𝑜𝑛𝑖_. The initial complex *C*_0𝑖_ can either unbind with rate 𝑘_𝑜𝑓𝑓𝑖_ or it can undergo 𝑁 sequential modifications to form activated complex *C*_𝑁𝑖_. Additionally, we assume that at any state the phosphorylated complex *C*_j𝑖_ can disintegrate with rate 𝑘_𝑜𝑓𝑓𝑖_ that reset complex to molecular form of receptor-ligand following the kinetic proof-reading scheme.

The time-dependent concentrations of the complexes *C*_0𝑖_, *C*_1𝑖_, … . . *C*_𝑁𝑖_ within the signaling pathway are described by the following Ordinary differential equations (ODE):

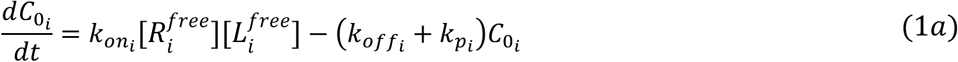

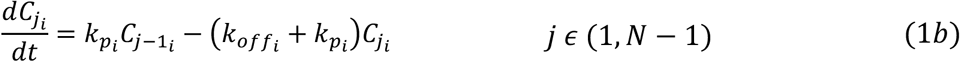

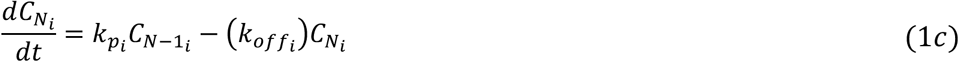

which lead to the following solutions at the steady-state limit

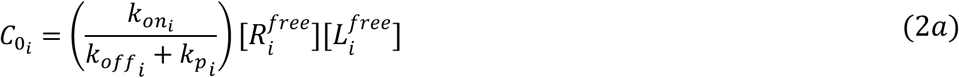

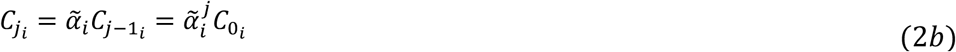

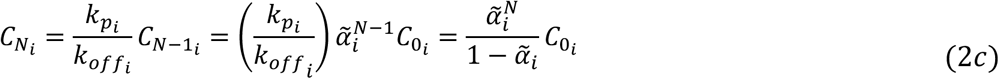

where 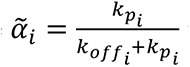 is the fraction of the concentration of intermediate complex going in forward direction before it disintegrates. The concentrations of free receptors and ligands in the steady state are 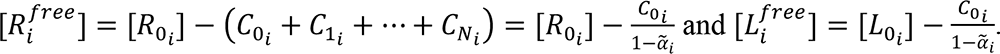. Substituting these into Eqn. (2a) lead to the following solution of *C_0i_* in the steady state

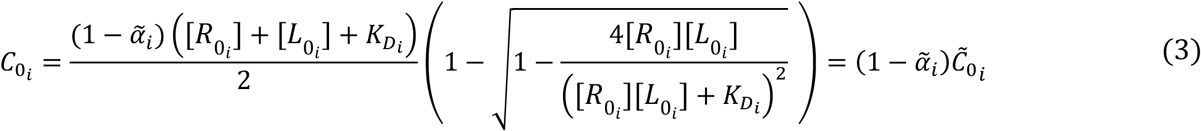

where the dissociation constant 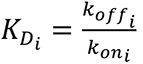. Thus, in the steady state

we get the solution of the final product using Eqns. (2c) and (3),

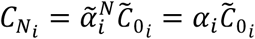

*Subcellular Scale (Stage-II):* We have assumed that the reversible inter-conversion between *V*𝑎𝑣1 (𝑊) and 𝑝*V*𝑎𝑣1 (𝑊^∗^) occurs by the kinase and phosphatase 𝐸_1_and 𝐸_2_, respectively (see following scheme).

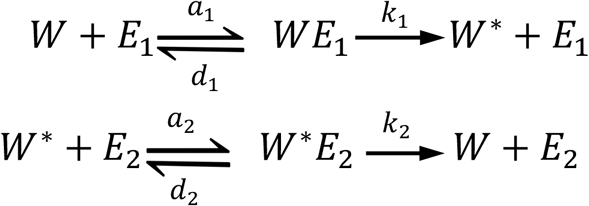

Enzyme 𝐸_1_ is the sum of the final activated complexes generated from CD33CAR, activating and adhesive receptor-ligand interactions (i.e., 𝐸_1_ = *C*_𝑁1_ + *C*_𝑁2_ + *C*_𝑁3_) and the enzyme 𝐸_2_ is the final activated complex of inhibitory receptor-ligand interaction (i.e., 𝐸_2_ = *C*_𝑁4_).

The time-dependent coupled ODE representing the kinetics of systems are:

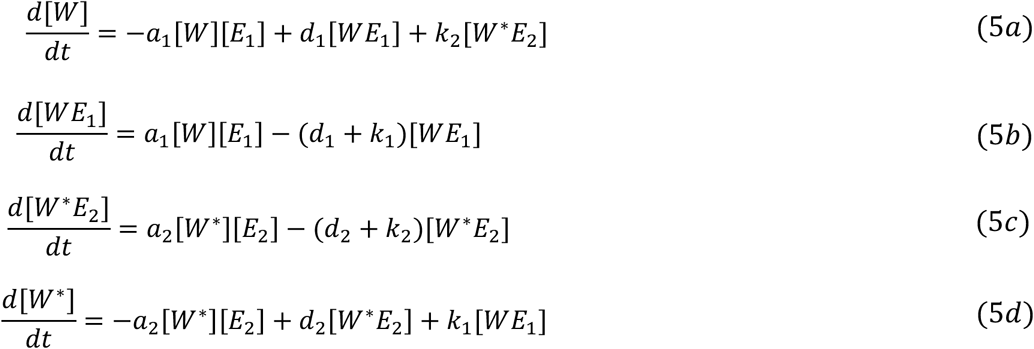

where [𝑊], [𝑊^∗^], [𝐸_1_], [𝐸_2_], [𝑊𝐸_1_] and [𝑊^∗^𝐸_2_] are the concentrations of Vav1, pVav1, kinase, phosphatase, intermediate complexes (in the phosphorylation and dephosphorylation reactions), respectively. At any time, the total concentration of Vav1 (unphosphorylated or phosphorylated or complex forms) is conserved 𝑊_𝑇_ = [𝑊] + [𝑊^∗^] + [𝑊𝐸_1_] + [𝑊^∗^𝐸_2_]. In the steady state, we get the solution of the fraction of pVav1 compared to total Vav1 as following (56)

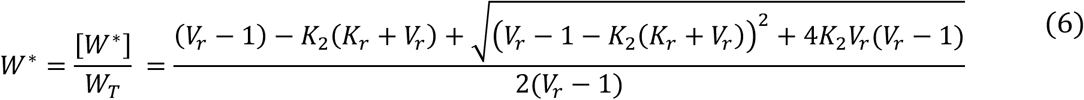

where 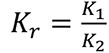 is constructed with the scaled Michaelis constant 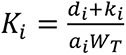 and 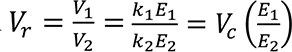

*Subcellular Scale (Stage-III):* The formation of pVav1 leads to the activation of the NK cell, which, when interacting with target cells, results in their lysis over time. Our model assumed the proliferation of the target cells with time. At any time *t*, the rate of change in the number of target cells having the ligands 𝐿 ≡ (𝐿_1_, 𝐿_2_, 𝐿_3_, 𝐿_4_) is:

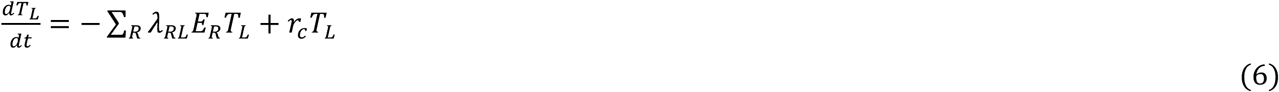

where E_R_ is the number of effector cells (NK cells) having receptors *R* ≡ {*R*_1_, *R*_2_, *R*_3_, *R*_4_} (with indices 1-4 corresponding to the types of receptors or ligands used in the model). The first term on the right-hand side is the loss term due to the interactions between target cells (𝑇_𝐿_) and NK cells (𝐸_*R*_) which leads to the lysis of the target cell with the proportionality constant *λ*_*R*𝐿_ determined by the distributions of the receptors and ligands. Our model assumed the constant *λ*_*R*𝐻_ is proportional to the concentration of pVav1 in the steady state, represented as *λ*_*R*𝐻_ = *λ*_c_𝑊^∗^with a lysis constant *λ*_c_. The summation is considered over different NK cells expressing various receptor types and quantities. The limits of the sum are decided by the number of the receptors or ligands based on their distributions on NK cells or the target cell. The second term is the gain in target cell 𝑇_𝐿_ due to its proliferation with 𝐿 ligands, where *r*_c_ is proliferation rate assumed to be constant.

Given the initial conditions {𝑇(0), 𝐸_*R*_(0)}, we solved Eq. (6) for given parameters 𝜃 ≡ 𝜃(*α*_1_, *α*_2_, *α*_3_, *α*_4_, *V*_c_, *K*_1_, *K*_2_, *λ*_c_, *r*_c_). We used the molecular distributions of three receptors (CD33CAR, LFA-1, and iKIR) for CAR-NK cells and the corresponding ligands (CD33, ICAM, MHC-1) for target cells as input. We numerically solved Eq. (6) using the function *solve_ivp* with ‘RK45’ method from the Python library *scipy.integrate* and applied the following formula to calculate the percentage of target cell lysis at time *t*

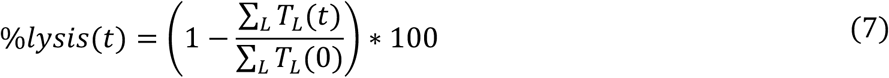

In the following, we discuss the methods that we have used to estimate parameters for different tumor cell lines.

**Parameter estimation:** We trained our model to estimate the parameters (*α*_1_, *α*_2_, *α*_3_, *α*_4_, *V*_c_, *K*_1_, *K*_2_, *λ*_c_, *r*_c_). We minimized the cost function for both CAR-NK and WT-NK simultaneously to find the optimal parameter values.

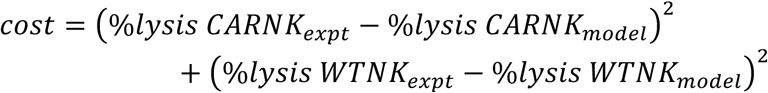

Minimization of cost function was performed using *least_squares* function of *scipy (a python module)* that uses “*Trust Region Reflective algorithm*”. The ranges of parameters for the search space are presented in Table 1.

***Maximum likelihood estimation:*** The CD33CAR expressions on CAR-NK cells showed a bimodal distribution (Fig. S3A). We modeled this using the superposition of two log-normal distributions 𝑝_1_(𝜇_1_, 𝜎_1_) and 𝑝_2_(𝜇_2_, 𝜎_2_), where mean (𝜇_1_, 𝜇_2_) and sigma (𝜎_1_, 𝜎_2_) are corresponding to respective log-normal distribution. We used a linear combination of these log-normal distributions: (1 − Ω) ∗ 𝑝_1_(𝜇_1_, 𝜎_1_) +Ω ∗ 𝑝_2_(𝜇_2_, 𝜎_2_) with weights (1 − Ω) and Ω for WT and CD33CAR-NK, respectively. We estimated five parameters (𝜇_1_, 𝜎_1_, 𝜇_2_, 𝜎_2_, Ω) that maximize the likelihood. We used a function *least squares* from *scipy (a python module)* that uses “*Trust Region Reflective algorithm*”.

𝑷𝒂𝒓𝒆𝒕𝒐 **optimization:** For targeted cancer immunotherapies like CAR-NK cell therapy there is a tradeoff between maximizing lysis of tumor cells while minimizing lysis of healthy cells. For example, CAR-NK cell can recognize and lyse healthy cells which express the targeted cognate antigen (i.e. CD33). However, healthy cells will express high amounts of major histocompatibility complex proteins that inhibit NK cell lysis. We set up a multi-objective optimization task to explore the signaling and receptor expression parameters in our model and calculate a Pareto front. This Pareto front represents the set of optimal parameters values where no objective can be further maximized without sacrificing the other. In our Pareto optimization scheme, we set two conflicting objectives: 1) minimization of healthy lysis and 2) maximization of tumor lysis.

We utilize the Non-dominated Sorting Genetic Algorithm II (NSGA-II) (44), a population-based method, for Pareto optimization to obtain the non-dominated solutions. This approach allows us to efficiently explore and identify optimal trade-offs among the above two objectives. We call functions *NSGAII* and *Problem* from *platypus* python library. We explore the optimal front within possible range of controlled parameters (*α*_1_, *α*_4_, Ω).

---

**Figure S1.**
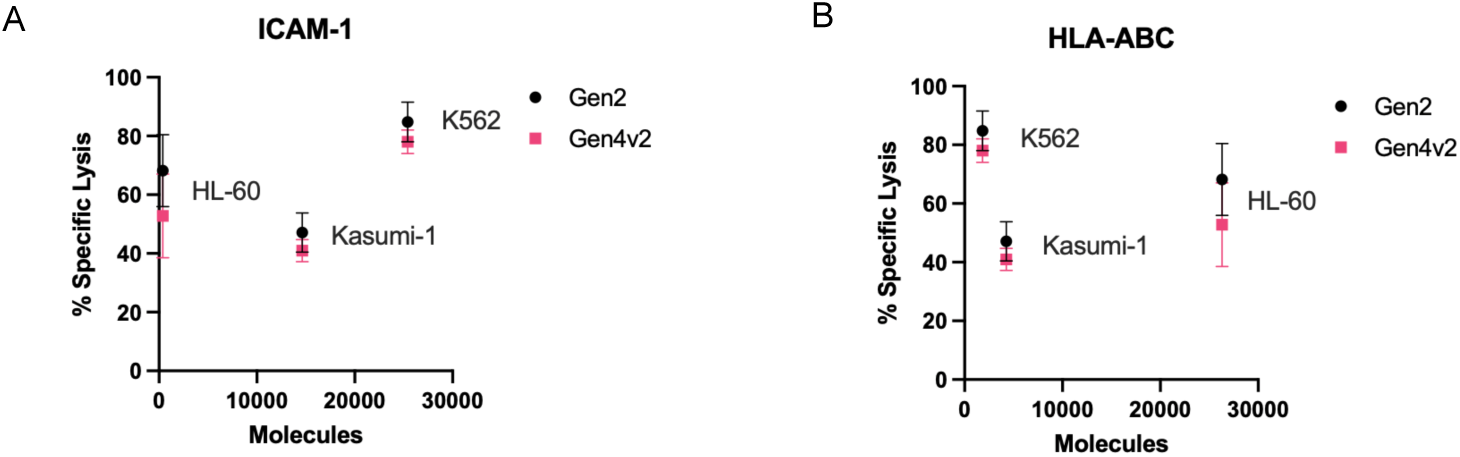
CAR-NK cytotoxicity displays non-monotonicity with ICAM-1 and HLA-ABC expression on target cells. A. Non-linear relationship CAR-NK cytotoxicity against K562, Kasumi-1, and HL-60 cell lines and ICAM-1 expression on target cells. B. Non-linear relationship CAR-NK cytotoxicity against K562, Kasumi-1, and HL-60 cell lines and HLA-ABC expression on target cells. ICAM-1 and HLA-ABC expression was quantified by flow cytometry. N = 3 donors.

**Figure S2.**
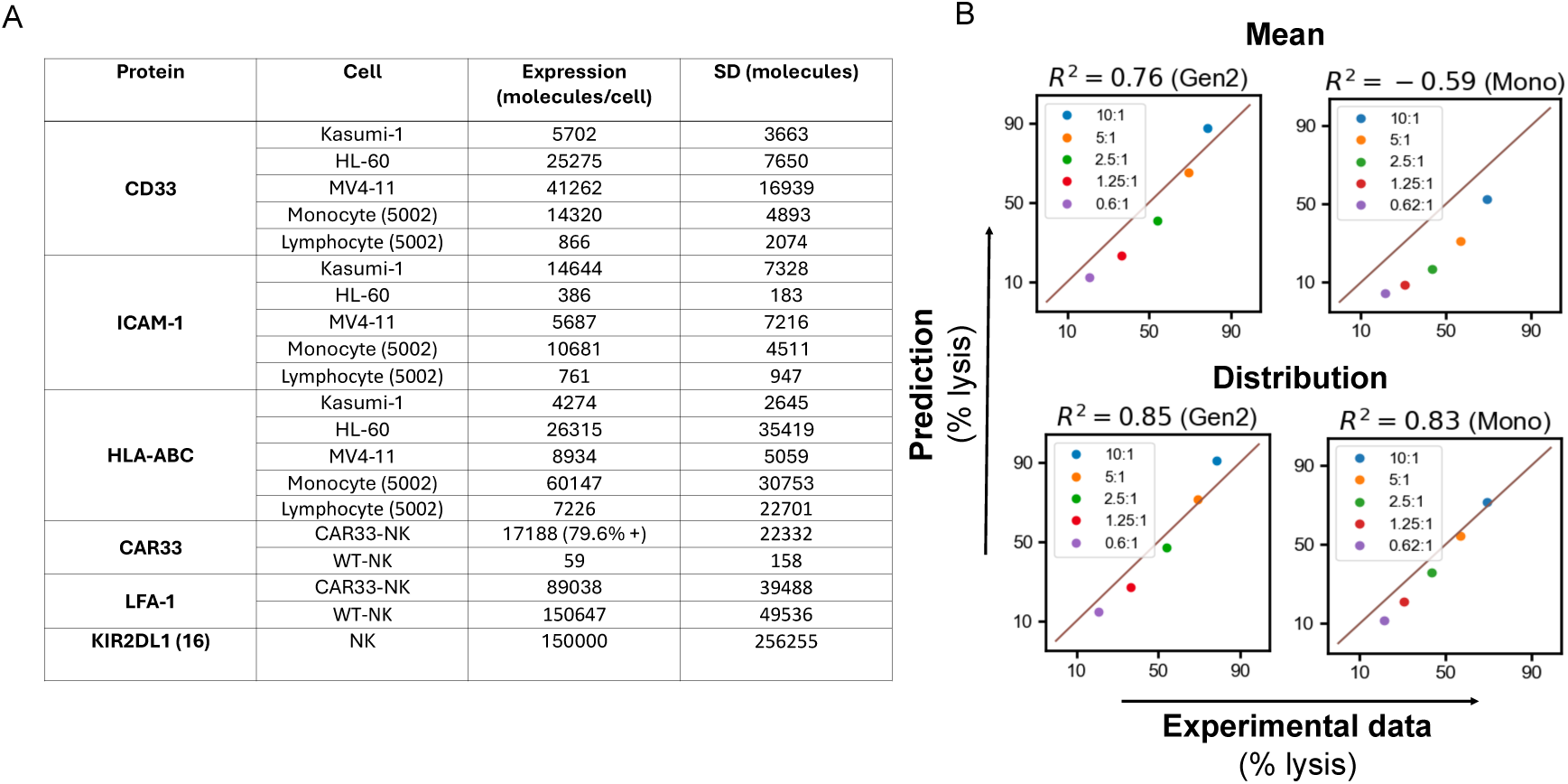
Ligand and receptor distributions. A. Table quantifying the molecular abundances of CD33, ICAM-1, HLA-ABC on Kasumi-1, HL-60, MV4-11, monocyte, and lymphocytes and CD33CAR, LFA-1, and KIR2DL1 expression on NK cells. Abundances quantified by quantitative flow cytometry. B. Comparison of model prediction accuracy using mean of ligand and receptor expressions (top) or distributions of ligand and receptor expressions (bottom).

**Figure S3.**
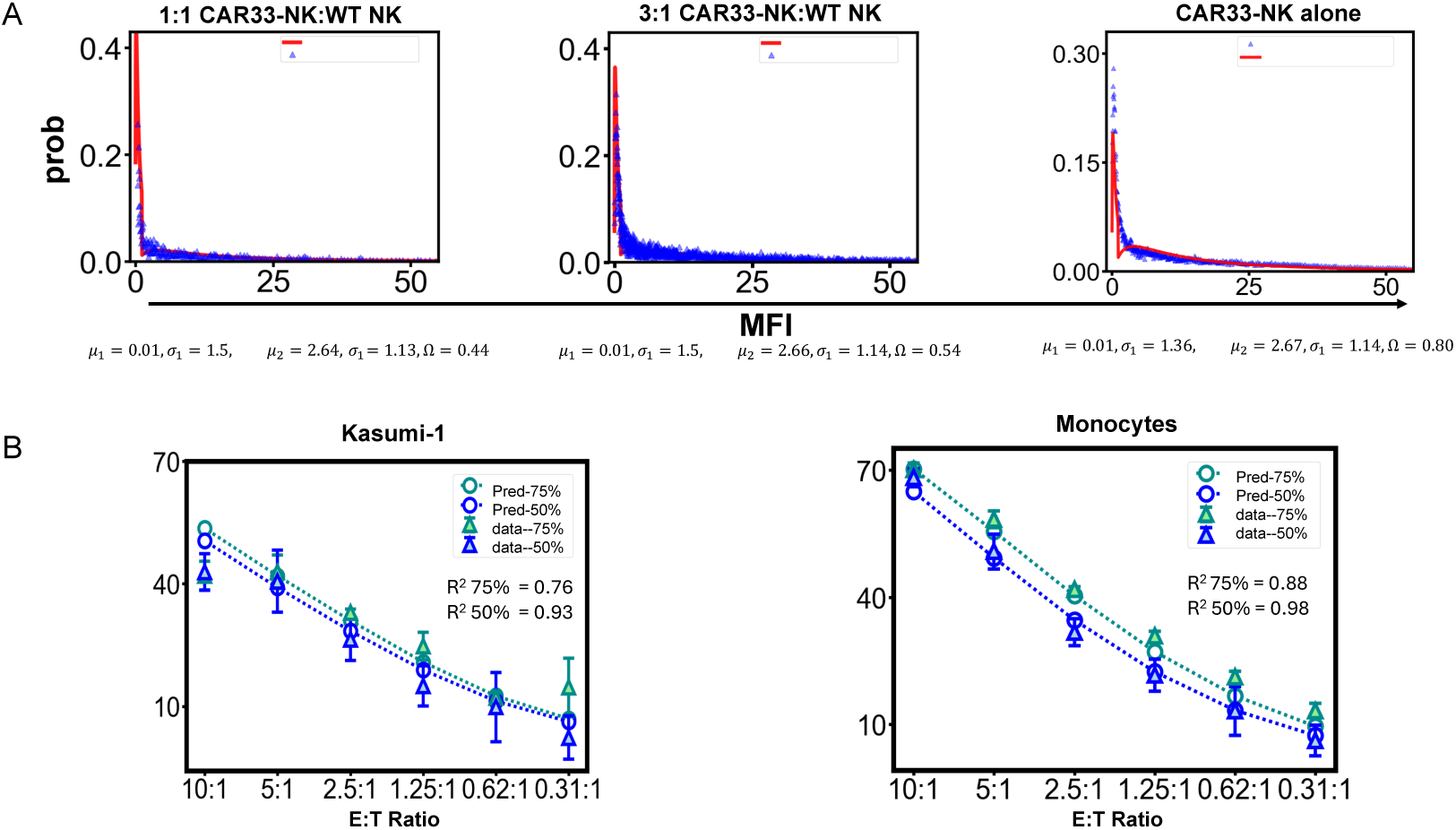
Maximum-likelihood estimation and model fitting. A. Maximum-likelihood estimation for mixtures of WT and CD33CAR-NK. We show parameter estimation for 50%, 75% and 100% CD33CAR where bule points represent the data and red line corresponds to superimposition of two log-normal distribution (1 − Ω)p_1_(𝜇_1_, 𝜎_1_) + Ω𝑝_2_ (𝜇_2_, 𝜎_2_). B. Model fitting and prediction of cytotoxicity for Kasumi-1 and monocytes with 50% CD33CAR and 75% CD33CAR. Mixtures of CD33CAR-NK and WT-NK were produced through sampling of experimental data.

## Acknowledgements

This work was supported by the Abigail Wexner Research Institute at Nationwide Children’s Hospital and the National Institutes of Health (NIH) (RO1AI146581-04) and the Department of Defense (CA200119). We acknowledge BioRender for generating schematic diagrams.

## Code availability

Data and code used in this paper are available at https://github.com/drsamalik/CAR-NK-cell-Modeling.

## Author Contributions

Conceptualization and study design: J.D., S.A., K.X. Data curation: K.X., K.A.B., M.N.K.

Statistical and computational analysis: S.A., H.R., I.N., W.C.S. Funding acquisition: J.D., D.A.L

Supervision: J.D., D.A.L., M.N.K. Visualization: S.A., K.X.

Writing – original draft: K.X., S.A., I.N., J.D. Writing – reviewing and editing: all authors.

## Data Availability Statement

Data are available from the corresponding authors upon request, except for identifying donor information.

